# Antigen-Specific modulation of Chronic Experimental Autoimmune Encephalomyelitis in Humanized Mice by TCR-like Antibody Targeting Auto-Reactive T-Cell Epitope

**DOI:** 10.1101/2024.07.15.603580

**Authors:** Ilana Goor, Efrat Altman, Inbar Arman, Maya Haus-Cohen, Yoram Reiter

**Affiliations:** Laboratory of Molecular Immunology and Immunotherapy, Faculty of Biology, Technion-Israel Institute of Technology, Haifa 3200001, Israel

**Keywords:** Antibodies, TCR-like antibody, MS/EAE, immunotherapy

## Abstract

The development and application of human T-cell receptor (TCR)-like antibodies (TCRL) recognizing disease-specific peptide-MHC complexes may prove an important tool for basic research and therapeutic applications.

Multiple Sclerosis is characterized by aberrant CD4 T cell response to self-antigens presented by class II MHC molecules. This led us to select a panel of TCRL Abs targeting the immunodominant autoantigenic epitope MOG35-55 derived from Myelin Oligodendrocyte Glycoprotein (MOG) presented on HLA-DR2 which is associated with Multiple Sclerosis (MS).

We demonstrate that these TCRL Abs bind with high specificity to human HLA-DR2/ MOG35-55 derived MHC class II molecules and can detect APCs that naturally present the MS-associated autoantigen in humanized EAE transgenic mouse model. The TCRLs can block ex vivo and in vivo CD4 T-cell proliferation in response to MOG35-55 stimulation in an antigen-specific manner. Most significant, administration of TCRL to MOG35-55 induced EAE model in HLA-DR2 transgenic mice both prevents and regresses established EAE. TCRL function was associated with reduction of autoreactive pathogenic T cells infiltration into the CNS, along with modulation of activated CD11b+ macrophages/microglia APCs.

Collectively, these findings demonstrate the combined action of TCRL Abs in blocking TCR-MHC interactions and modulating APC presentation and activation, leading to a profound antigen-specific inhibitory effect on the neuroinflammatory process, resulting in regression of EAE.

Our study constitutes an in vivo proof-of-concept for the utility of TCR-like antibodies as antigen-specific immunomodulators for CD4-mediated autoimmune diseases such as multiple sclerosis (MS), validating the importance of the TCR-MHC axis as a therapeutic target for various autoimmune and inflammatory diseases.

## Introduction

Multiple sclerosis (MS) is a chronic neuroinflammatory autoimmune disease of the central nervous system (CNS) characterized by immune cell infiltration across the blood-brain barrier (1–5). This infiltration promotes inflammation, demyelination, gliosis, and neuroaxonal degeneration, disrupting neuronal function. Early in lesion formation, autoreactive CD4 T lymphocytes play a crucial role by mounting aberrant responses against CNS autoantigens. These autoreactive CD4 T cells are activated by innate immune cells, including dendritic cells, natural killer (NK) cells, macrophages in the periphery, and microglia, astrocytes, and infiltrating monocytes in the CNS (1–5). These cells mediate damage through antigen presentation on MHC class II molecules, presenting CNS autoantigens to autoreactive T cells. The strong genetic association of MS with the HLA-DRB1*15:01 allele underscores the importance of MHC class II in the disease’s pathogenesis (6,7).

Current immunomodulatory drugs for MS effectively reduce immune cell activity but have systemic effects and are often associated with significant side effects, such as flu-like symptoms and progressive multifocal leukoencephalopathy. Therefore, there is an unmet need for more specific therapies that can eliminate autoreactive immune responses without broadly compromising the immune system (8).

Decades of clinical and basic experimental research into multiple sclerosis point to distinct immunological pathways that drive disease relapses and progression. The variation in clinical manifestations of the disease correlates with the spatiotemporal dissemination of lesioned sites of pathology within the central nervous system (CNS). These lesions are a hallmark of multiple sclerosis and are caused by immune cell infiltration across the blood–brain barrier (BBB) that promotes inflammation, demyelination, gliosis and neuroaxonal degeneration, leading to disruption of neuronal functions (1–5). T cells appear early in lesion formation, and the disease is considered to be autoimmune, initiated by autoreactive lymphocytes that mount aberrant responses against CNS autoantigens. Intervention of the infiltration of immune cells from the periphery into the CNS has been the main target of currently available therapies for multiple sclerosis. Although these broad-spectrum immunomodulatory drugs reduce immune cell activity and entry into the CNS and decrease relapse frequency, they are often associated with side effects. These range from flu-like symptoms and the development of other autoimmune disorders to malignancies and even fatal opportunistic infections such as progressive multifocal leukoencephalopathy (8). Thus, more specific therapeutic targets that can be efficaciously modulated without inducing such significant adverse reactions are required.

The immune cells which characteristically infiltrate from the periphery into the CNS in MS, must be specifically activated in order to induce the associated tissue damage. During the establishment of central tolerance in the thymus, most autoreactive T cells are deleted; however, this process is imperfect, and some autoreactive T cells are released into the periphery. In the healthy population, peripheral tolerance mechanisms keep these cells in check. If this tolerance is decreased, by reduced function of regulatory T (T-Reg) cells and/or the increased resistance of effector B cells and T cells to suppressive mechanisms specific autoreactive B cells and T cells can be activated in the periphery to become aggressive effector cells. This activation may be in the form of molecular mimicry, novel autoantigen presentation, recognition of sequestered CNS antigen released into the periphery or bystander activation. Genetic and environmental factors, including infectious agents and cigarette smoke constituents, contribute to these events. Once activated, differentiated CD4+ T helper 1 (TH1) and TH17 cells, CD8+ T cells, B cells and innate immune cells can infiltrate the CNS, leading to inflammation and tissue damage. B cells trafficking out of the CNS can undergo affinity maturation in the lymph nodes before re-entering the target organ and promoting further damage. In the common experimental model of MS; experimental autoimmune encephalomyelitis (EAE) infiltrating CD4+ T cells are re-activated in the CNS by antigen-presenting cells (APCs), including CD11c+ dendritic cells (DCs), with the resulting inflammatory response leading to monocyte recruitment into the CNS, as well as naive CD4+ T cell activation through epitope spreading that further fuels the inflammation (2–5). Autoantigen presentation by APCs to CD4+ T cells is thus the epicentre of disease initiation and progression and the major histocompatibility complex:T cell receptor (MHC:TCR) interaction between the APCs and the pathogenic autoreactive T cells is the central driving force of immune activation.

This autoantigen presentation of class II HLA-DR molecules to the pathogenic autoreactive TCR on T cells is thus a most specific check-point in the inflammatory process. We therefore aimed to block these specific MHC: TCR interactions.

The principle of our experimental design is to generate a T-cell receptor-like antibody (TCRL) that mimics TCR specificity binding to human MHC class II complexes presenting a myelin-derived autoantigen peptide on APCs (9–12).

In the past such a TCRL antibody was selected towards HLA-DR2 molecules complexed with an immunodominant myelin basic protein (MBP) peptide (residues 85-99) and was used for visualization of MBP T cell epitopes in multiple sclerosis lesions (9). Therapeutic activity of TCRL antibodies *in vivo* for the prevention or treatment of established experimental autoimmune encephalomyelitis (EAE) was never demonstrated.

Previous work in our lab had shown that the anti HLA-DR4/GAD_555-567_ TCRL antibody, directed towards a peptide derived from type-1-diabetes autoantigen glutamic acid decarboxylase (GAD), was able to block the T cell response *in vivo* (10).

Herein we present the first *in vivo* proof of concept that a TCRL antibody can cause significant regression of an established autoimmune disorder in a humanized transgenic mouse model of EAE. We demonstrate our working hypothesis that inducing tolerance to myelin antigens would have an effect on EAE initiation and progression and show that the TCRL antibody exerts its biological activity through a combined action of blocking MHC: TCR interactions between the APCs and pathogenic autoreactive CD4+ T cells as well as modulating specifically CD11b+ APCs that present the autoantigen.

## Results

### Selection and characterization of TCR-like antibodies that recognize MOG-specific HLA-DR2 restricted autoreactive T cell epitope

In our experimental setup through the presented studies, we have used *in vitro* and *in-vivo* systems of human HLA-DR2 APCs as well as a humanized mouse model of EAE, in which immunization of HLA-DR2-transgenic mice with the MS-associated autoantigen Myelin Oligodendrocyte Glycoprotein (MOG)-derived mouse peptide mMOG_35-55_ results in a chronic EAE (13).

In order to isolate TCRL antibodies specific to HLA-DR2/mMOG_35-55_ we first expressed and purified recombinant HLA-DR2 molecules using constructs in which the intracellular domains of the DR-A1∗0101 and DR-B1∗1501 chains were replaced by leucine-zipper dimerization domains for heterodimer assembly (supplementary data Figure 1A, B). Purified recombinant DR2 complexes were loaded with the mMOG peptide or myelin basic protein (MBP)-derived peptide as a control.

Using the purified recombinant DR2/mMOG_35-55_ complexes, we screened a large human naïve Fab antibody phage display library (14) and isolated in several screening campaigns several TCR-like antibodies with variable affinities termed 2G10, I4Z, and I2A all of which recognized specifically recombinant DR2/mMOG_35-55_ complexes (Figure 1A). The Fab antibody constructs were transformed into a full chimeric human-mouse IgG antibody constructs (IgG2a) by fusing the Fab variable domains to the scaffold of murine IgG2a Fc. The chimeric TCRL Abs were expressed in Expi293 cells and its TCR-like specificity was further tested on recombinant HLA-DR2 complexes displaying mMOG_35-55_ or control MBP peptide as well as native complexes presented on HLA-DR2-positive B cell line (MGAR) loaded with the mMOG_35-55_ peptide as observed by microscopy (Figure 1B) and flow cytometry (Figure 1C). The TCRLs did not bind to recombinant complexes (Figure 1A) or to MGAR B cells that present the control MBP peptide (Figure 1C). The proper loading of these peptides to the B cell APCs was confirmed with a conformational sensitive anti-HLA-DR2 antibody (Figure 1C). Dose dependent binding of the TCRLs exhibited saturable and specific binding with apparent EC50 of high to medium affinity (10-100 nM) (Figure 1C) supplemented with SPR data that confirmed these affinities (supplementary Figure 1C).

**Figure 1:**
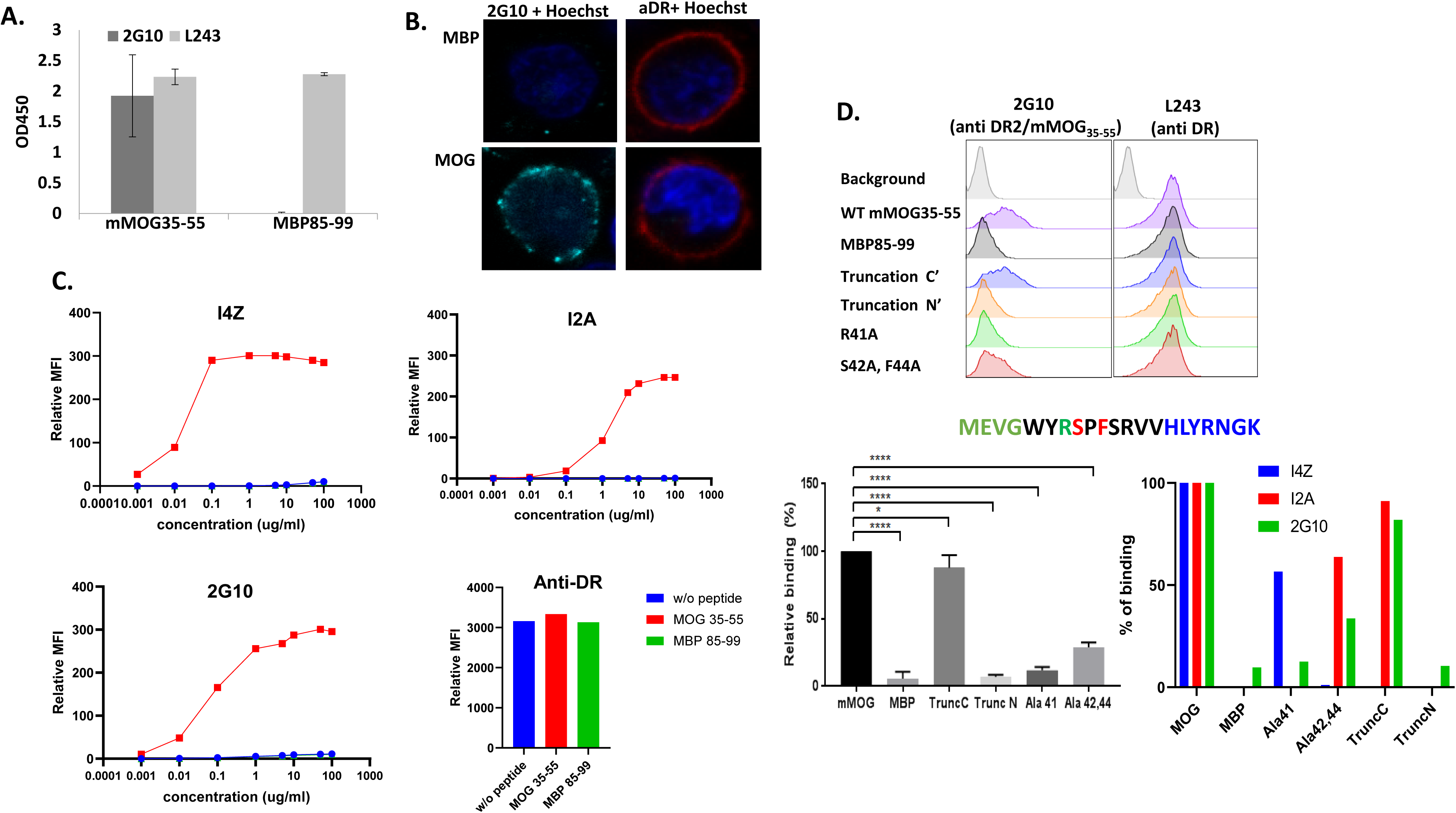
Isolation and characterization of anti-MOG/DR2 TCRL Abs. A) Binding specificity of 2G10 TCRL Ab. ELISA binding assay with 2G10 TCRL Ab towards recombinant HLA-DR2 complexes presenting mMOG_35-55_ but not MBP_85-99_ peptides. Anti-DR served as additional control. B) Immunofluorescence staining of DR2+ MGAR cell line loaded with mMOG_35-55_ (Right) or MBP_85-99_ (Left) peptides. Cells were stained with anti-HLA-DR (red) and 2G10 TCRL (green). C) MGAR cells were loaded with MOG, MBP or no-peptide. Cells were tested for mIgG2a I4Z, I2A or 2G10 staining at different antibody concentrations (0.001-100 ug\ml). Anti-DR served as positive control. D) MGAR cells were loaded with altered MOG 35-55 peptides, in addition to original MOG 35-55 and MBP 85-99. Cells were tested for I4Z, I2A or 2G10 staining. Top - representative flow cytometry analysis. Amino acid sequence of the wild type (wt) mMOG_35-55_ peptide. Lower panel - average of three independent experiments examining 2G10 binding (Right) or 2G10, I4Z and I2A (Left) normalized to the binding of the WT peptide (100%) and presented as percent relative binding. Right: Statistical analysis was performed by one way ANOVA, Dunnett’s multiple comparisons test.

The high specificity of the TCRL Abs, similar to TCR, was validated by their ability to distinguish minor changes in the mMOG_35-55_ peptide sequence. As shown in Figure 1D, we tested the ability of the TCRLs to recognize DR2-positive APCs loaded with a set of mMOG_35-55_ altered peptide ligands: R41A, S42A+F44A, as well as N-terminal and C-terminal truncated mMOG (mMOG_39-55_ and mMOG_35-48_, respectively). These mutations did not significantly affect peptide binding to recombinant HLA-DR2 as judged both by competition assays using biotinylated wild-type MOG peptide and the 4 altered non-biotinylated peptides (not shown). Truncation at the C- terminus of the peptide did not affect the binding intensity of the I2A and 2G10 Abs compared to their binding to the wild type DR2/mMOG_35-55_ but abolished the binding of I4Z. In contrast, the double Alanine substitutions at position 42 and 44 had a significant effect on 2G10 TCRL binding (∼70% reduction in binding) and distinct variable effect on I4Z and I2A. Moreover, Alanine substitution at position 41 (R41A) affected binding of 2G10 and I2A while truncation at the N-terminus of the peptide had detrimental effect and completely (>95%) abolished binding of all three TCRL Abs (Figure 1D). It is important to note that the TCRL Abs do not bind to empty recombinant HLA-DR2 molecules (not shown). These results suggest that TCRL Abs bind the DR2/mMOG_35-55_ epitope in a peptide-sequence dependent manner utilizing multiple specific contact residues which mimic TCR-like fine specificity features.

### TCR-like antibodies can detect EAE-disease-specific MOG-derived/HLA-DR2 peptide complexes on the surface of APCs

Next, we evaluated the ability of the TCRL Abs to recognize APCs that present HLA- native DR2/MOG complexes, by APC peptide pulsing or naturally-occurring antigen processing and presentation. TCRL Abs (shown for 2G10) were able to bind specifically all splenocytes-derived APCs subpopulations (B cells, dendritic cells and macrophages) pulsed with the mMOG35-55 peptide (Figure 2A). Upon confirming the specificity of the Abs, we tested whether they can recognize and detect APCs that present mMOG35-55 peptide in the context of class II HLA-DR2 molecules after naturally-occurring antigen processing and presentation in the context of mMOG35- 55-induced EAE murine model using HLA-DR2 Tg mice (15). Sixteen days post EAE induction we used TCRL Abs to stain APCs by flow cytometry and detect HLA- DR2/mMOG35-55 complexes on the cell surface, investigating which APCs present the autoantigen in spleens and spinal cords harvested from mMOG35-55 -immunized mice. Although, B cells are the most abundant APC population in the spleen, we did not detect presentation of mMOG35-55 peptide on these cells (Figure 2B) but were able to detect significant presentation of mMOG35-55 on the CD11c+ dendritic cells subpopulation in comparison with a healthy mice control as expected by their role in antigen uptake (Figure 2B). These TCRL stained CD11c+ dendritic cells constituted a significant (2-4%) number of the total CD11c+ cells in the spleen of EAE mice compared to controls. Macrophages exhibited some degree of mMOG35-55 presentation which upon statistical analysis was not significant. In contrast to the relatively low frequency of presentation in the periphery, we found substantial presentation of mMOG35-55 in the spinal cord of EAE-diseased mice compared with healthy controls, both in the microglia (tissue resident macrophages) subpopulation (CD11b+/CD45^low^) and in the activated macrophage (CD11b+/CD45^high^) (Figure 2C, D). Although both CD11b+ subpopulations present at high frequency (10-40%) mMOG35-55, we demonstrate that the frequency of presentation of mMOG35-55 by CD11b+/ CD45^low^ cells is higher compared to the CD11b+/ CD45^high^ APC subpopulation. Staining data observed for the I4Z and I2A TCRLs detecting CD11b+/CD45^high/low^ cells was variable (Figure 2F) in mild and severe EAE but significant compared to healthy mice or HLA-DR2 negative mice.

**Figure 2:**
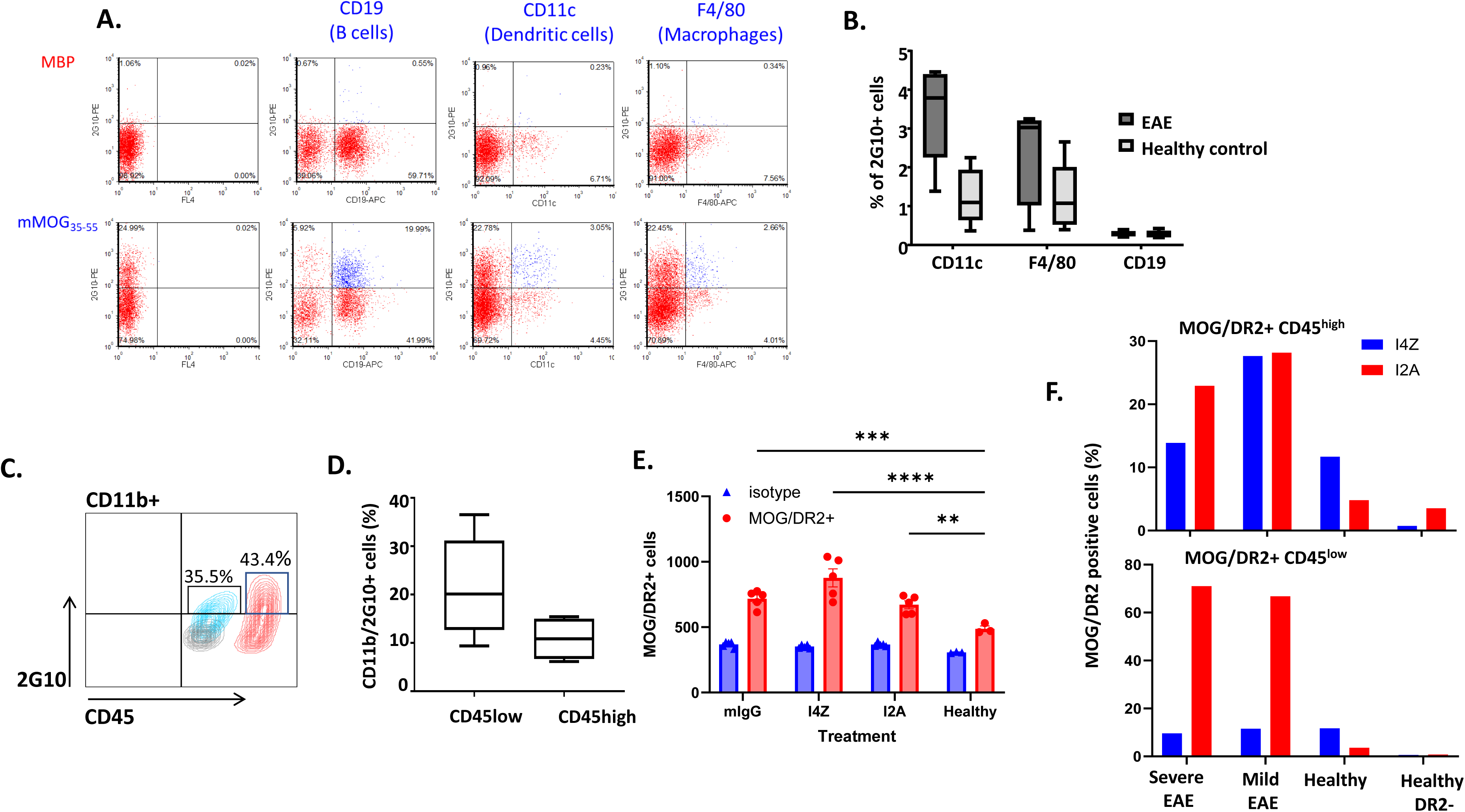
MOG/DR2 presentation can be detected by anti-MOG/DR2 TCRL Abs within various tissues. A) Binding of 2G10 TCRL Ab to various APCs derived from splenocytes of HLA-DR2 Tg mice by flow cytometry. APCs were pulsed with the mMOG35-55 and control MBP85-99 peptides and subsequently stained with 2G10 TCRL Ab and APC markers CD19, F4/80, CD11b. B) Average percentage of splenetic DCs (CD11c), macrophages (F4/80) and B cells (CD19) presenting HLA-DR2/mMOG_35-55_ as detected by 2G10 TCRL Ab in EAE-diseased or healthy controls. C) Representative staining of CD11b+ cells in the spinal cord. Phenotype of DR2/ mMOG_35-55_ CD11b+ cells in the spinal cord or EAE-diseased mice. Average percentage of spinal cord CD11b+/CD45^high^ and CD11b+/CD45^low^ that present HLA-DR2/mMOG_35-55_ in EAE mice. E) MOG/DR2 presentation by CD45low monocytes within the CNS of TCRL-treated and control-treated EAE-diseased mice. Healthy DR2 mice served as control. Presentation was assessed by FACS at experiment endpoint. Statistical analysis was performed by one way ANOVA, Dunnett’s multiple comparisons test. F) MOG/DR2 presentation by CD45high or CD45low monocytes within the CNS of severely or mildly EAE-diseased mice. Healthy DR2 mice and Healthy C57bl/6 DR2(-) mice served as control. Presentation was assessed by FACS at experiment endpoint.

Investigations with these TCRL Abs enabled the first direct detection of myelin-derived autoantigen presentation in the CNS and in the periphery during EAE. This approach also identified the APCs involved in autoantigen presentation, as well as the interplay between the high frequency of presentation at the site of inflammation and the leakage of presentation in the periphery.

### The HLA-DR2/MOG-specific TCRL antibodies are functional *in vitro* and can mediate inhibition of T cell proliferation and induce antibody ADCC effector functions

Next, we examined the biological activity properties of the TCRL Abs with the following functions tested: (i) ability to specifically block stimulation of autoreactive T cells as measured by inhibition of proliferation and/or cytokine release and (ii) mediated antibody effector function in the form of ADCC. Figure 3 shows these assays and demonstrate that TCRL Abs are functional and can mediate these functions in vitro. Inhibition of proliferation in an ex-vivo assay is shown for 2G10 (Figure 3A) in which DR2-Tg mice or DR2/MBP-TcR double Tg mice were immunized with mMOG35-55 or MBP85-99, respectively. Ten days post immunization a recall T-cell proliferation assay was performed with immunized mice-derived CFSE-labelled splenocytes that were re-stimulated with the priming peptide (mMOG35-55 or MBP85-99) in the presence of increasing concentrations of 2G10 TCRL Ab. As shown in Figure 1A, 2G10 TCRL Ab inhibited specifically mMOG35-55 reactive CD4 T cell proliferation in a dose dependent manner but did not affect the proliferation of MBP85-99-reactive T cells, demonstrating ex-vivo that the 2G10 TCRL Ab can specifically block MHC: TCR interactions and potentially mediate antigen-specific immunomodulation of T cell activation/proliferation. Inhibition of T cell activation/cytokine release is also shown in Figure 3B and C for I4Z and I2A, demonstrating peptide titration effect on T cell stimulation and inhibition by TCRL, as well as specificity (Figure 3C) in which no T cell stimulation was observed with MBP peptide. Further, specific ADCC functions mediated by the TCRLs were demonstrated as shown in Figure 3D-F. 2G10 (Fig. D) I4Z and I2A (Fig. E and F) exhibited specific ADCC function towards MOG-presenting DR2 cells which was dose dependent and comparable to a pan anti-DR or anti-CD20 antibody. We also utilized an ADCC dead mutant version of the TCRLs (D265A mutation (16)) and demonstrated that this version cannot mediate ADCC (Fig. 3D, F).

**Figure 3:**
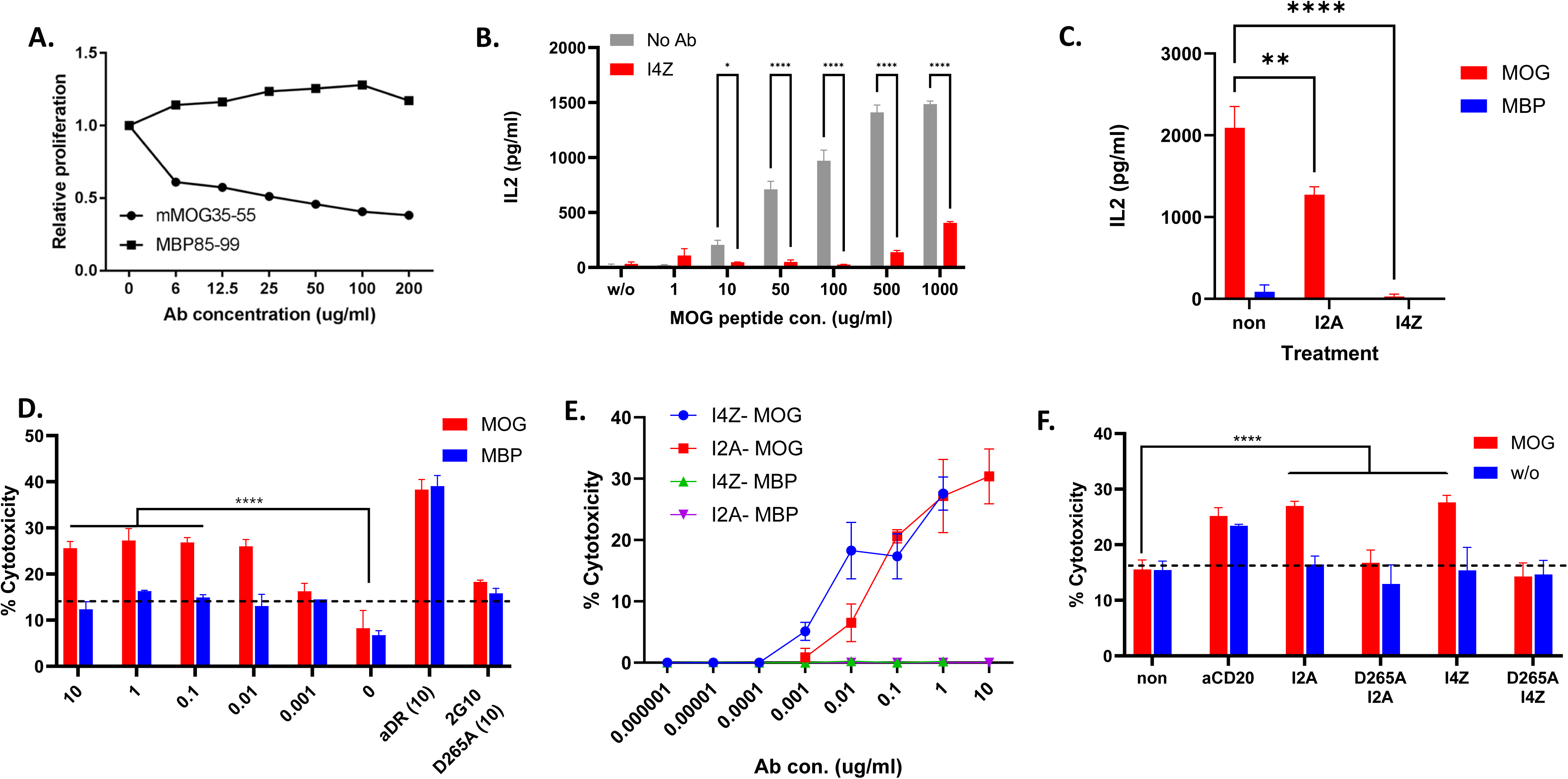
HLA-DR2/MOG-specific TCRL antibodies are functional *in vitro* and can mediate inhibition of T cell proliferation and induce ADCC effector functions. A) Antigen specific inhibition of MOG-specific T cell proliferation. HLA-DR2-Tg mice were immunized with mMOG_35-55_/CFA and HLA-DR2/MBP-TCR-Tg mice were immunized with MBP_85-99_/CFA. 10 days post immunization, CFSE-labelled splenocytes were re-stimulated with mMOG_35-55_ or MBP_85-99_ peptide, respectively, in the presence of increasing concentrations of 2G10 and 5 days later CD4 T cell proliferation was measured. Results were normalized in comparison to proliferation without 2G10 TCRL Ab (Relative proliferation). B, C) MGAR cells were loaded with various concentrations of MOG peptide and incubated as described with 10 ug/ml of I4Z (B and C) or I2A (C). After 2h MOG-directed CAR-T cells were added at 1:1 E:T ratio for 24h and IL-2 secretion was assessed. D-F) MGAR DR2+ APCs were loaded with MOG or MBP peptides and tested for specific killing by 2G10 (D), I4Z and I2A (E and F) TCRL Abs. ^35^S-methaionine radioactive or LDH release assays were used to access cytotoxicity with human PBMCs (E) or murine NK cells (D and F) or at 10:1 E:T ratio. D265A ADCC - dead Mutant TCRL constructs and anti-CD20 Abs were used as controls.

Altogether, our data demonstrate that the TCRL Abs can mediate antigen-specific modulation and functions toward APCs and T cells exerting the two postulated modes of actions; blocking T cell functions by competition with TCRs through binding of the TCRL Ab to prevent stimulation/activation of pathogenic T cells and effector function through antigen-specific ADCC on APCs that present the autoantigen.

### TCRL Abs prevent EAE in HLA-DR2 humanized murine model

As a first step in examining the potential of the TCRL Abs as a therapeutic modulatory agent, we tested their ability to prevent mMOG_35-55_ induced EAE in the DR2 Tg mice. In this EAE prevention experiment, EAE was induced by immunization with mMOG_35- 55_ in CFA and Pertussis toxin (Ptx). On day 0 and 2 post immunization mice were treated with 200µg 2G10 mIgG, a control mIgG or Vehicle (PBS) (I.P. injection). As shown in Figure 4, the TCRLs were able to ameliorate EAE severity, with the 2G10 TCRL Ab (Fig 4C) having the most significant effect in decreasing EAE scores in comparison to the control groups. I2A was also functional with statistical significance of several independent experiments (Fig. 3 A, B). 13 days post immunization, when the control group reached the peak of disease with an average disease score of 2.1 and 2.2 in the mIgG or PBS controls respectively, the majority (12/16; 75%) of 2G10 treated mice had an EAE score ≤ 0.5. 3/16 (19%) 2G10 treated mice had an EAE score of 1 and only one of the 2G10 treated mice did not respond to the TCRL Ab treatment and developed a severe disease (EAE score=3) (Figure 4D).

**Figure 4:**
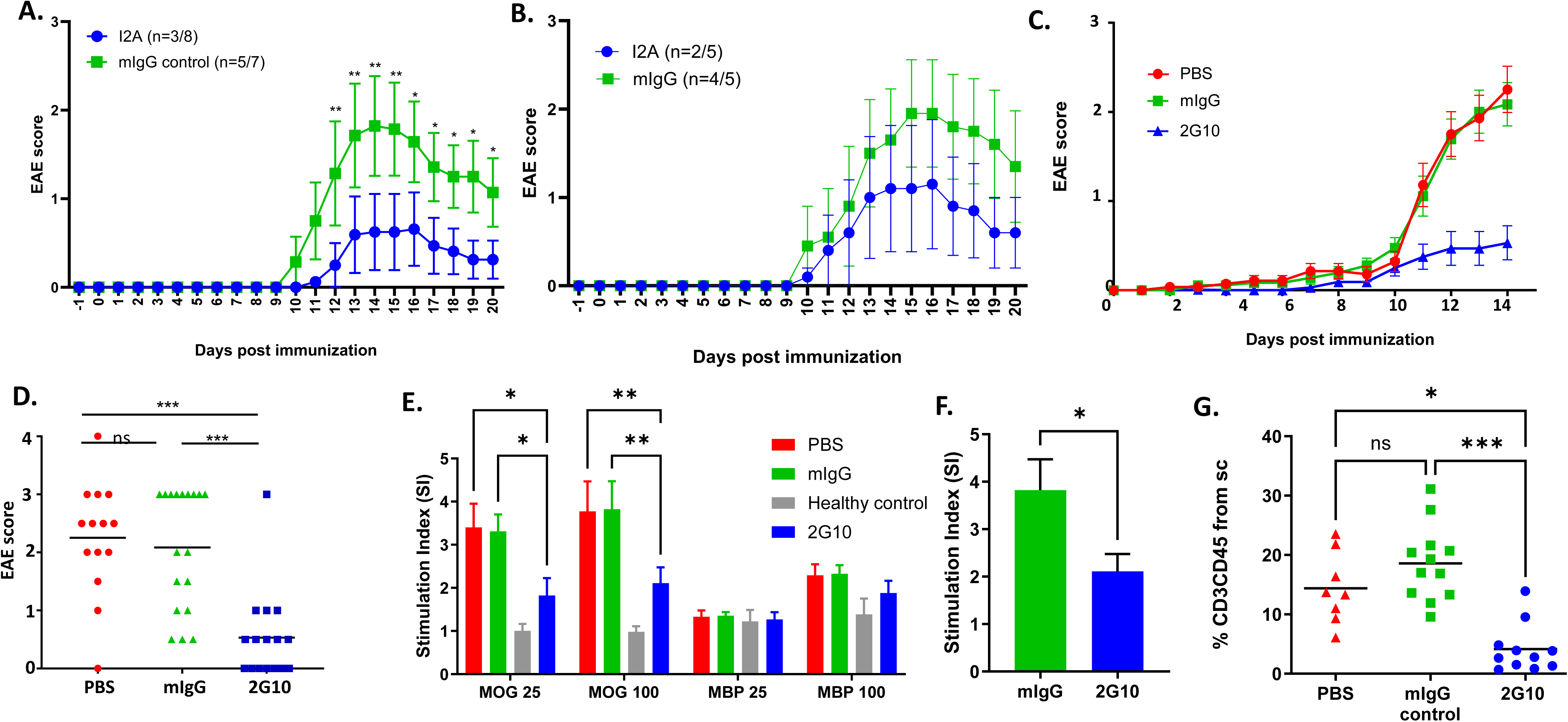
TCRL Abs prevent EAE in HLA-DR2 humanized murine model. EAE was induced in DR2 male mice by MOG _35-55_/CFA immunization. Mice were treated i.p at days -1, 2, 5 and 8 with I2A (A, B) or at days 0 and 2 with 2G10 (C) TCRL Ab. mIgG or PBS served as control. A-C) Changes in EAE score in response to I2A or mIgG (A, B), or 2G10, PBS or mIgG (C). D) Individual EAE scores at day 14 post immunization of C. Changes in EAE score of individual mice in response to 2G10, PBS and mIgG control. E, F) *Ex-vivo* T cells (E) and CD4 isolated T cells (F) proliferation in spleen of EAE treated mice. Splenocytes from 2G10 treated and controls were re-stimulated with the mMOG_35-55_ priming epitope or a control MBP_85-99_ peptide (25µg/ml or 100µg/ml) and proliferation was monitored by [^3^H]-thymidine uptake. Data was normalized to proliferation without peptide and is presented as stimulation index (SI). (G) CD3+/CD45+ cells in spinal cords collected from a subset of the mice in C determined by flow cytometry at the peak of disease (day 14). 2G10, mIgG control and PBS. In all assays data was compared using Kruskal-Wallis followed by a Dunn’s multiple comparison test or two-way ANOVA followed by Dunnett’s multiple comparison test. Error bars, s.e.m.

In an attempt to study the mechanism of action of the TCRL Ab and to test whether the prevention of EAE by 2G10 was facilitated by blocking *in-vivo* priming of mMOG_35-55_ specific T cells, we examined the recall response to the mMOG_35-55_ peptide of splenocytes derived from 2G10 treated mice versus the mIgG control or PBS treated mice. At day 14-post mMOG_35-55_ peptide immunization, splenocytes were harvested and re-stimulated with the priming mMOG_35-55_ peptide or the MBP_85-99_ peptide. As shown in Figure 4E, splenocytes from mice treated with the 2G10 TCRL Ab exhibited a significantly lower mMOG_35-55_-specific proliferation compared to mIgG control or vehicle-treated mice. Recall proliferation assays with splenocytes primed with the MBP_85-99_ peptide did not show significant changes in stimulation between TCRL treated and control mice (Figure 4E).

The effect on proliferation was governed mostly by CD4 T cells as the inhibitory effect induced by the 2G10 TCRL Ab was profound when CD4 was isolated prior to re-stimulation with the mMOG_35-55_ peptide (Figure 4F). We also examined whether the ability of 2G10 TCRL Ab to inhibit *in-vivo* T cell priming and proliferation is antigen specific, by performing re-call assays on splenocytes isolated from influenza hemagglutinin-derived (HA) peptide- immunized mice treated with the 2G10 TCRL Ab or vehicle only. In contrast to the significant blockade of mMOG35-55-reactive T cell priming; we did not observe a difference in HA peptide-mediated proliferation in the 2G10 TCRL Ab treated versus untreated mice (not shown). These results indicate that *in-vivo* blocking and inhibition of T cell priming mediated by the 2G10 TCRL Ab is antigen specific.

Overall, results are in line with our proposed mechanism of action and demonstrate the ability of the 2G10 TCRL Ab to block and inhibit *in-vivo* T cell priming and proliferation and consequently prevent disease.

Antigen specificity is thus setting the basis of our experimental design to validate the TCRL Ab as a therapeutic modality that can be effective in eliminating or modulating self-response without generally compromising the immune system. To further investigate the biological consequences of 2G10 TCRL Ab treatment in the EAE prevention model we examined the migration of T cells and APCs to the CNS. Spinal cord examination of 2G10 TCRL Ab treated mice revealed a significantly lower level of infiltrating T cells in comparison to the mIgG treated control group indicating that blocking and inhibition of T cell priming also inhibited or prevented spinal cord inflammation through reduced T cell migration into the CNS (Figure 4G). The 2G10 TCRL Ab had no influence in the EAE prevention model on the frequency of CD11b+/CD45^high^ and CD11b+/CD45^low^ APCs in the CNS (not shown). Interestingly, we were able to detect infiltrating CD11b+/CD45^high^ APCs as early as 8 days post immunization, before the appearance of EAE clinical signs, suggesting that the immunization allowed activated macrophages to cross the blood brain barrier without additional stimulation/feedback and that the 2G10 TCRL Ab was unable to prevent that effect.

### TCRL Abs modulate established EAE in HLA-DR2 humanized murine model

Antigen specific TCRL Ab prevention of EAE by blocking T cell priming, activation and proliferation at the MHC: TCR axis can be useful in the therapeutic window between relapses of MS. However, treatment of an established and ongoing EAE disease has greater implications. To test the ability of the TCRL Abs to treat established EAE, we induced chronic EAE in DR2-Tg mice, and begun treatment when clinical score was ≥ 2, usually occurring usually 10-12 days post immunization. EAE mice were treated with 2G10 TCRL Ab at day 12 and 14 post immunization and were followed for disease signs. As shown in Figure 5A-D, mice injected with the TCRL Abs had variable responses in this model with 2G10 having a significant improvement effect of EAE clinical signs as early as 24h after the first injection. A representative established EAE treatment experiment over time is shown in Figure 5A and a summary of 5 independent established EAE treatment experiments with the 2G10 TCRL Ab is shown in Figure 3C, D (day 0 refers to the first dose of 2G10 TCRL when clinical score is >2, 10-12 days post immunization). While the control mice reached an average clinical score of 3 at the peak of disease, the 2G10 treated mice improved gradually, until they reach an average clinical score of 1 (Figure 5A, C). The cumulative disease index (CDI) of individual treated mice was also significantly lower in 2G10 treated mice compared to the CDI of the control mice (Figure 5D). The significantly improved clinical signs in mice treated with the 2G10 TCRL Ab, (i.e., disease score of 1 or even lower) were stable for >50 days post treatment (not shown). Interestingly, the modulatory effects of I2A and I4Z were not significant and I4Z even enhanced disease scores. These differential effects can be due to multiple factors such as optimal affinity or specific epitope binding as shown for peptide-dependent binding in TCRL characterization.

**Figure 5:**
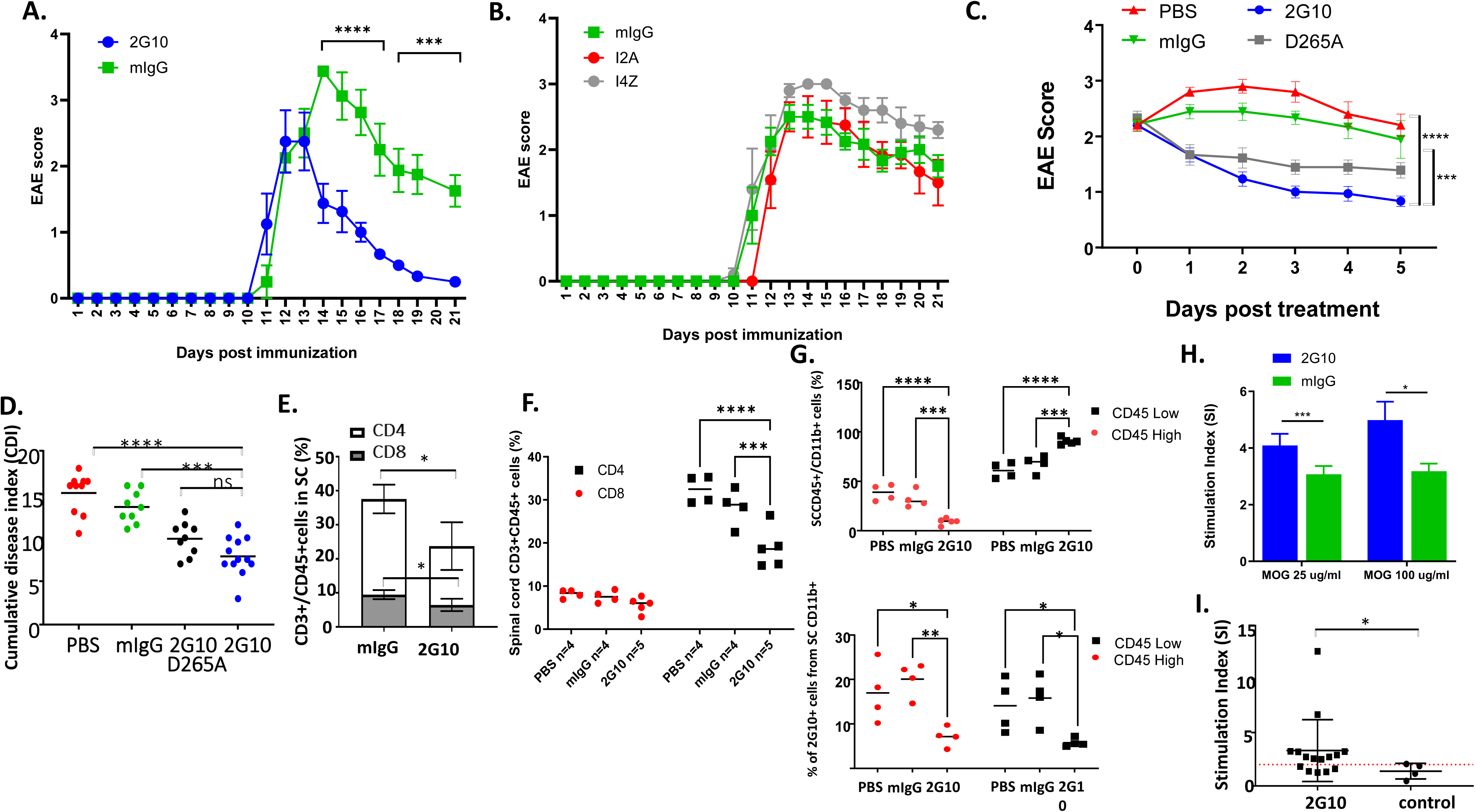
TCRL Abs modulate established EAE in HLA-DR2 humanized murine model. EAE was induced in DR2 male mice by MOG 35-55/CFA immunization. At disease score ≥ 2 mice were treated i.p with 200µg Ab/mice, following a second dose after two days. A-C) Changes in EAE score in response to 2G10 and mIgG (A), I4Z, I2A or mIgG (B) or 2G10, PBS, mIgG control and 2G10 D265A (C). D) individual EAE score of the mice in C. Data presented as cumulative disease index (sum of the daily scores since the immunization). E, F) Percentage of CD3+CD45+ cells. G) Top: Percentage of mononuclear cells in spinal cords collected from a subset of the mice in C determined by flow cytometry 5 days after treatment initiation. Bottom: Percentage of CD11b+CD45^high^ and CD11b+CD45^low^ cells presenting the mMOG35-55 peptide. Data was compared using Mann-Whitney U test or Kruskal-Wallis followed by a Dunn’s multiple comparison test or two-way ANOVA followed by Dunnett’s multiple comparison test. Error bars, s.e.m. (H) T cell proliferation in 2G10 TCRL Ab treated and control EAE mice. *Ex-vivo* proliferative response to re-stimulation with the mMOG35-55 priming or control MBP85-99 peptide determined by [3H]-thymidine uptake in spleens collected from a subset of the mice 5 days after the initiation of treatment. Data was normalized to proliferation without peptide and is presented as stimulation index (SI). (I) T cell proliferation towards MBP in 2G10 TCRL Ab MOG-induced EAE treated and healthy mice. In vitro proliferative response to re-stimulation with MBP85-99 peptide determined by [3H]-thymidine uptake in spleens collected from a subset of the mice 5 days after the initiation of treatment. Data was normalized to proliferation without peptide and is presented as stimulation index (SI). In all assays data was compared using Mann-Whitney U test or Kruskal-Wallis followed by a Dunn’s multiple comparison test or two-way ANOVA followed by Dunnett’s multiple comparison test. Error bars, SEM.

A single aspartic acid to alanine mutation in the Fc domain has been shown to almost completely abolish the binding of IgG2a and IgG2b to all four classes of Fc gamma receptors and the activation of the complement (12). This mutation was introduced into 2G10 TCRL Ab and as shown in Figure 5C, D, the mutant D265A 2G10 TCRL Ab was less efficient that the wild type TCRL Ab in treating established EAE, although the differences were not significant. These results may suggest that most of the 2G10 TCRL Ab biological mode of action is related to its ability to inhibit T cell priming, activation and proliferation through blocking the MHC:TCR interactions and the pathogenic T-cell:APC axis of interactions. Although the possibility that TCRL Ab activity stems from Fc-mediated ADCC and CDC biological activity cannot be excluded.

Next, we measured and compared the infiltration of lymphocytes and active macrophages to the CNS (spinal cord) in EAE treated vs. untreated and control mice to determine the influence of 2G10 TCRL Ab on the migration of these cells. As shown in Figure 5E, F, 2G10 treated mice had a significantly lower percentage of T cells in the spinal cord. 2G10 TCRL Ab treatment effected both CD4+ T cells, which were the majority of the T cells in the spinal cord, and on CD8+ T cells. Not only did the 2G10 TCRL Ab have a significant effect on T cells frequency in the CNS, it also exhibited a significant influence on APCs in the CNS. As shown in Figure 5G (and supplementary Figure 3 for independent experiments), 2G10 TCRL Ab led to a highly significant reduction in the frequency of CD11b+/CD45high activated macrophages in the spinal cord and restored the composition of a healthy spinal cord with a predominant CD11b+/CD45low population of tissue resident microglial APCs.

Another indication of reduced inflammation in the spinal cord following treatment with 2G10 TCRL Ab is the reduction in the frequency of APCs that present the autoantigen, i.e. the mMOG35-55 peptide. As shown in Figure 3G, the reduction in mMOG35-55 peptide presenting APCs was observed for both subpopulations of CD11b+/D45high and CD11b+/CD45low.

Altogether, these results indicate that TCRL Ab-mediated blockade of pathogenic T cell re-activation is sufficient to suppress an ongoing chronic EAE. Moreover, the suppression of T-cell reactivation induced a broad effect on spinal cord inflammation and thus resulted in decreased infiltration of activated APCs into the CNS as well as significant reduction in Myelin presentation on top of the direct effects on T cell functions.

In contrast to the reduction in the frequency of T cells in the spinal cord after treatment with the 2G10 TCRL Ab we observed an accumulation of mMOG35-55 reactive T cells in the spleens of treated mice as demonstrated by the significantly higher mMOG35- 55-specific proliferation in splenocytes harvested from 2G10 TCRL Ab treated mice compared to vehicle-treated mice (Figure 5H). These results indicate that in 2G10 TCRL Ab treated mice mMOG35-55 specific T cells are maybe sequestered in the spleen and upon the inhibitory effect on their reactivation and proliferative capacity as well as their ability to migrate to the CNS.

Interestingly, stimulation of splenocytes from 2G10 TCRL Ab treated mice with MBP peptide, 16-18 days post EAE induction through immunization with mMOG35-55 peptide resulted in higher proliferation in comparison to healthy mice (Figure 5I). This indicates that the inhibitory therapeutic activity of the 2G10 TCRL Ab can be exerted even in the presence of MBP reactive T cells.

In summary, the results presented herein suggest that antigen-specific targeting of the TCR: MHC axis has potent therapeutic potential and lays grounds for the development of TCRL Ab- based immunotherapeutic agents. These TCRL Abs can be developed for antigen-specific therapy of MS, and could potentially be employed to other autoantigens implicated in various T-cell mediated autoimmune diseases.

Targeting disease associated APCs that process and present myelin epitopes have the advantage of overcoming the dynamic T cell repertoire in autoimmune diseases. Moreover, macrophages and myeloid cells at the site of inflammation effect the disease course by releasing oxygen species and therefore their targeting may have beneficial effects that are not restricted to their role in T cell activation. The direct visualization of MOG presenting APCs at the site of inflammation as shown in this study for the disease EAE model makes them a validated target.

## Discussion

The experiments presented in this study demonstrate for the first time an *in vivo* proof of concept for targeting class II MHC-peptide complexes that present an autoantigen on the surface of APCs that are involved in the activation of pathogenic autoreactive T cells in autoimmunity. Such mode of targeting was facilitated by a TCRL Ab that is directed toward the highly characterized MS-associated HLA-DR2-restricted MOG_35-55_ epitope which was shown to induce severe chronic EAE in HLA-DRA/DRB1*1501-Tg mice (HLA-DR2).

We have shown that administration of 2G10 TCRL Ab to MOG_35-55_ induced EAE in HLA-DR2 transgenic mice prevented the disease and most significantly was able to ameliorate established EAE in these mice leading to a robust and durable inhibition and reversal of disease in animal models.

Elucidating the mode-of-action of 2G10 revealed that it inhibits the activation of HLA- DR2/mMOG_35-55_ reactive T-cells *in-vivo* and reduces the infiltration of autoreactive pathogenic T lymphocytes into the CNS. Moreover, modulation of activated CD11b+ macrophages/microglia APCs that present the mMOG_35-55_ in the CNS was also demonstrated. Collectively our data demonstrates the combined action of the TCRL in blocking TCR-MHC interactions as well as modulating APC presentation and activation leading to a profound antigen-specific inhibitory effect on the neuroinflammatory process and regression of EAE.

In previous studies, we have applied our TCRL Ab approach to generate and characterized TCRL Abs against auto-reactive T-cell epitopes associated with type-1- diabetes (T1D) and demonstrated that such TCRL Abs are able to block *in vivo* re-stimulation of CD4 T cells recognizing the T1D associated autoantigen GAD in the context of class II HLA-DR4 molecules (10,40). However, the TCRL Ab approach and targeting of autoantigen associated class II HLA-DR epitopes was never tested in an animal model of autoimmune disease as performed herein for chronic EAE.

The TCRL Ab was also used to detect the specific APCs that present the targeted autoantigen in the spleen and CNS of EAE mice and thus is a crucial tool for elucidating the mode of action of the TCRL Ab.

The long-term implications of these studies introduce the scientific basis for further evaluation of this approach in pre-clinical humanized transgenic mouse models of EAE to further demonstrate that the TCRL Ab approach can induce antigen-specific tolerance as a novel immunotherapy for MS and as a proof of concept for other autoimmune and inflammatory diseases.

Class II autoantigen specific TCRL Abs could confer clinical benefits through three potential modes of action: (i) blocking TCR-APC interactions at the core disease axis of TCR:MHC (ii) elimination of autoantigen presenting APCs which present not only the targeted epitope or autoantigen-derived class II MHC-peptide complex but also other autoantigens using a highly functional TCRL Ab with ADCC and CDC effector functions, or (iii) immunomodulation of APCs and their microenvironment using the TCRL Ab as a vehicle to deliver an immunosuppressive cytokine (immunocytokine), thus conferring an immunosuppressive environment at the site of inflammation leading to suppression or inhibition of the immune response towards the auto-antigen (17,18, 33–37).

TCR-like antibodies have shown potential as TCR mimicking molecules in research and therapeutic development for autoimmune diseases due to their high affinity and ease of engineering. This approach is being developed also in line with studies employing engineered soluble TCRs (19–23). TCR-like antibodies have been shown to identify specific populations of antigen-presenting cells (APCs) in autoimmune conditions. For example, in multiple sclerosis (MS) (9) TCRL antibodies have identified microglia macrophages as predominant autoreactive APCs in MS lesions. Similarly, in rheumatoid arthritis (RA), TCRL antibodies specific for the human cartilage glycoprotein (HC gp-39) epitope presented by the HLA-DR4 allele have identified dendritic cells presenting this autoantigen in inflamed joints of RA patients (39). In celiac disease, TCRL antibodies have identified plasma cells as the primary cells presenting gluten peptides (31,32).

The ’armed’ TCRL Ab approach can target, using single antibody specificity towards a well-defined and characterized autoantigen, APCs that present multiple autoantigens and thus have a profound effect on disease progression. TCRL Ab-induced elimination of autoantigen- specific APCs is expected to suppress the trigger and response against all postulated autoantigens thus changing the course and progression of the disease and may even present a potential solution for antigen spreading.

Despite their potential, TCRL antibodies have not been extensively explored as therapeutics for autoimmune diseases. However, preclinical studies in cancer have demonstrated their therapeutic potential, particularly in targeting intracellular tumor antigens presented by MHC-I molecules (11–12,25–30). A limitation in cancer therapy is the low coverage of TCRL antibodies per cell due to MHC-I downregulation on tumors. In contrast, MHC-II molecules are typically upregulated on auto-APCs in autoimmune diseases, providing a more robust target for TCRL antibodies.

In line with the above-mentioned modes of actions, therapeutic strategies for autoimmune diseases utilizing TCRL antibodies could involve:

Depletion of pathology-driving cells, similar to cancer therapy, but with a focus on reestablishing immune balance rather than broad cell elimination. For instance, anti- CD20 antibodies deplete B cells in RA and MS but come with concerns such as long-term benefits and side effects. The use of bispecific Antibodies (BsAbs) which could combine the specificity of anti-CD20 and anti-autoantigen p/MHC for targeted depletion of pathology-related B cells is of great potential. Non-Depleting TCRL mAbs can modulate immune responses by limiting autoantigen-MHC accessibility as shown in this study, thus reducing the activation of cognate T cells. This could be achieved through engineering antibodies with low Fc receptor binding. Autoimmune modulators coupled with TCRL mAbs like toxins, immunoregulatory cytokines (e.g., IL-10, TGF- β), or antibodies targeting inflammatory cytokines (e.g., TNF, IL-6, IL-1β) could be delivered to autoantigen-MHC-II enriched sites to reestablish immune tolerance. TCRL antibodies could be used in chimeric antigen receptor (CAR) formats to construct CAR T cells. In diabetic NOD mice, TCR-like antibodies and CAR T cells expressing an insulin peptide/MHC-II TCRL antibody modulated autoimmunity (41–44). In another approach, redirecting regulatory T cells (Treg) to the autoimmune milieu could suppress autoreactive effector T cells. Nanobodies with TCR-like specificity can be also developed for various therapeutic modalities (45,46).

Altogether, the data we present in this work indicates that the biological mode of TCRL Ab action is mostly related to the ability to inhibit T cell priming, activation and proliferation through blocking the MHC:TCR interactions and the pathogenic T-cell:APC axis of interactions. This inhibition leads to reduced migration to the inflammatory sites of pathogenic T cells and APCs that present the autoantigen. The possible role of the TCRL Ab through Fc-mediated ADCC and CDC biological activity cannot be excluded and needs to be further explored through optimization of TCRL Ab constructs and experimental systems.

Specifically for MS, further investigation of the neuro-inflammation process as a whole is required in order to better delineate which inflammatory and neurodegenerative mechanisms are truly distinct but occur in parallel and which are inextricably associated. This analysis will be crucial in aiding the design of more effective therapeutic strategies. A major goal for future treatment of multiple sclerosis may thus be the simultaneous, early targeting of peripheral immune cell function and of CNS-intrinsic inflammation, potentially through combinatorial therapies designed to effectively and specifically modulate these two immunological arms of the disease, along with the provision of neuroprotective or neurodegenerative drugs. The TCRL Ab approach which targets the disease core immune cell functions at the TCR: MHC axis may be a good starting point to develop antigen-specific modulatory agents that will suppress both peripheral immune functions as well as CNS-associated inflammatory processes.

## Materials and methods

### Production of recombinant HLA-DR2 in S2 cells

Construct, transfection, and expression of recombinant MHC class II have been previously [8]. Briefly, in these constructs, the intracellular domains of the DR-A1∗0101 and DR-B1∗1501 chains were replaced by leucine-zipper dimerization domains for heterodimer assembly. Bir A recognition sequence for bitionylation was introduced to the C-terminus of the DR-A1∗0101 chain. The constructs were cloned between the Bgl II and EcorI restriction sites into the pMT/BiP/V5-His vector. The vectors were co-transfected with pCoBlast selection vector into S2 cells using Fugene reagent (Promega). Stable single-cell clones were verified for protein expression. Upon induction of the cells with CuSO4, the supernatant was collected and DR2 complexes were affinity purified using anti-DR LB3.1 (ATCC number HB-298) mAb. The purified DR2 complexes were biotinylated by Bir-A ligase (Avidity) and characterized by SDS-PAGE. The biotinylated complexes were loaded with the mMOG35-55 peptide or the MBP85-99 for 48h at 37 in PBS pH7.4. The right folding of the complexes was verified by recognition of anti-DR conformation sensitive mAb (L243) in ELISA binding assay.

### Selection of phage Abs on biotinylated complexes

Selection of phage Abs on biotinylated complexes was performed as described before. Briefly, a large human Fab library containing 3.7 x 10_10_ different Fab clones was used for the selection (14). Phages were first pre-incubated with streptavidin-coated paramagnetic beads (200 µl; Dynabeads) to deplete the streptavidin binders. The remaining phages were subsequently used for panning with decreasing amounts of biotinylated MHC-peptide complexes. The streptavidin-depleted library was incubated in solution with soluble biotinylated DR2/mMOG35-55 complexes (20ug for the first round, and 5ug for the following rounds) for 30 min at room temperature. Streptavidin-coated magnetic beads (200 µl for the first round of selection and 100 µl for the following rounds) were added to the mixture and incubated for 10–15 min at room temperature. The beads were washed extensively 12 times with PBS/0.1% Tween 20 and an additional two washes were with PBS. Bound phages were eluted with triethylamine (100 mM, 5 min at room temperature), followed by neutralization with Tris-HCl (1 M, pH 7.4), and used to infect E. coli TG1 cells (OD = 0.5) for 30 min at 37°C.

### Expression and purification of soluble recombinant Fab Abs

TG1 or BL21 cells were grown to OD600 = 0.8–1.0 and induced to express the recombinant Fab Ab by the addition of isopropylthio-β-galactoside (IPTG) for 3–4 h at 30 °C. Periplasmic content was released using the B-PER solution (Pierce), which was applied onto a prewashed TALON column (Clontech). Bound Fabs were eluted using 100 mM imidazole in PBS. The eluted Fabs were dialyzed against PBS (overnight, 4°C) to remove residual imidazole.

### Construction of whole IgG Ab

The H and L Fab genes were cloned for expression as mouse IgG2a Ab into the eukaryotic expression vector pCMV/myc/ER. For the H chain, the multiple cloning site, the myc epitope tag, and the endoplasmic reticulum (ER) retention signal of pCMV/myc/ER were replaced by a cloning site containing recognition sites for BssHI and NheI followed by the mouse IgG2a constant H chain region cDNA isolated by RT-PCR from human lymphocyte total RNA. A similar construct was generated for the L chain. Each shuttle expression vector carries a different antibiotic resistance gene. Expression was facilitated by cotransfection of the two constructs into the Expi293 expression system (Thermo Fisher Scientific). The desired Ab were further purified from their supernatant using protein A affinity chromatography. SDS-PAGE analysis of the purified protein revealed homogenous, pure IgG with the expected molecular mass of ∼150 kDa.

### ELISA with purified Fab Abs

Binding specificity of individual soluble Fab fragments was determined by ELISA using biotinylated MHC/peptide complexes or peptides. ELISA plates (Falcon) were coated overnight with BSA-biotin (1 μg/well). After being washed, the plates were incubated (1 h at room temperature) with streptavidin (10 μg/ml), washed extensively, and further incubated (1 h at room temperature) with 5 μg/ml of MHC/peptide complexes or peptides. The plates were blocked for 30 min at room temperature with PBS/2% skim milk and were subsequently incubated for 1 h at room temperature with 5 μg/ml soluble purified Fab. After washing, plates were incubated with horseradish peroxidase-conjugated/anti-human-Fab antibody. Detection was performed using 3,3′,5,5′- tetramethylbenzidine (TMB; Sigma).

### Spinal Cord, Brain and splenocytes isolation

Spleens were removed from euthanized animals under sterile conditions and single cell suspensions of leukocytes were prepared by disaggregation of the tissue through a 100 μm nylon mesh (BD Falcon, Bedford, MA). Cells were washed once with RPMI 1640 supplemented with 10% heat-inactivated FBS (Biological Industries), then incubated with RBC Lysis buffer (Sigma) for 5 min to remove red cells, then washed in PBS and re suspended in culture media.

Brains and spinal cords were passed through 100 μm mesh screens and washed as above. Cells were resuspended in 80% Percoll (GE Healthcare, Pittsburgh, PA) then overlaid with 40% Percoll to establish a density gradient and centrifuged at 1600 rpm for 30 min following a method previously described [3]. Leukocytes were collected from the resultant interface, counted, and resuspended in FACS buffer (PBS -0.1% BSA), for further analysis.

### Flow cytometry

For exogenous loading of APCs: DR2-EBV-transformed B lymphoblast MGAR cell line (IHW#9104) was incubated for ON with cRPMI medium containing various concentrations of one of the following peptides, as mentioned: mMOG35-55 (MEVGWYRSPFSRVVHLYRNGK), mMOG trunc N (WYRSPFSRVVHLYRNGK), mMOG trunc C (MEVGWYRSPFSRVV), mMOG S42A, F44A (MEVGWYRAPASRVVHLYRNGK), mMOG R41A (MEVGWYASPFSRVVHLYRNGK) or MBP85-99 (ENPVVHFFKNIVTPRTP). All peptides were purchased from LifeTein at 95% purity. Cells were washed and incubated with various concentrations of TCRL Abs, or control aDR Ab, as mentioned, for 1 h at 4 °C, followed by incubation for 45 min at 4 °C with anti-mouse PE secondary Ab. Cells were then washed and analysed by FACS Calibur flow cytometer (BD).

Antibodies — leukocytes were labelled with a combination of the following antibodies obtained from BioLegend: CD4 APC/Cy7 (GK1.5), CD3 FITC (145-2C11), CD45 APC (30F-11), CD11b PE (M1/70), anti-human HLA-DR PE (L243), anti-mouse IgG PE (Poly4053), Strp –BV, 7AAD. eBiosceince: CD11c APC (N418), F4/80 APC (BM8). BD: CD19 APC (1D3), Jackson ImmunoResearch: anti-human IgG PE

Extracellular staining — single cell suspensions were washed and resuspended in staining buffer (PBS- 0.1% BSA). Fc receptors were blocked with TruStain fcX™ (anti- mouse CD16/32) (clone 93) and cells were incubated with either hIgG-TCRL Ab or biotin-TCRL, then washed and incubated with anti-human-BV or Strp-BV, respectively, and conjugated monoclonal antibodies (mAbs) listed above. Unbound mAbs were washed away with staining buffer and 7AAD was used in order to exclude dead cells prior to flow cytometry analysis using a BD LSR cytometer (BD Biosciences)

### Cytotoxicity assays

Antibody dependent cell cytotoxicity was measured using a commercially available LDH Non-Radiactive Cytotoxicity Assay (Promega) according to the manufacturer’s instructions. Target MGAR cells, loaded with 400 ug/ml MOG or MBP peptide, were incubated for 2h at the presence of 10 ug/ml mIgG2a TCRL Ab, mIgG2a-D265A TCRL Ab, anti-mouseCD20 and no Ab. After 2h all cells were washed and co-incubated with freshly isolated mouse NK cells (using EasySep Mouse NK Cell Isolation Kit (StemCell)) at 10:1 effector to target ratio for 4 hours, and the concentrations of LDH in supernatants was measured.

For methionine based killing assay target MGAR cell were labelled with S35- methionine radioactive isotope for 8-12 hours at 37°C and subsequently washed and loaded with 200ug/ml MOG or MBP peptide overnight. S35 labelled cells were incubated for 2h at the presence of different concentrations (102-10-6 ug/ml) of hIgG1 TCRL Ab. Anti-humanCD20 or no Ab served as control. After 2h all cells were washed and co-incubated with human total PBMCs at 20:1 effector to target ratio for 6 hours. Supernatants S35-methionine concentration was measured, as indication for cell death.

### Relative Cytotoxicity was calculated directly using the following equation

LDH concentration values were corrected by subtracting culture medium background, effector cells spontaneous release and target cells spontaneous release. The percent specific cytotoxicity was calculated as follows:

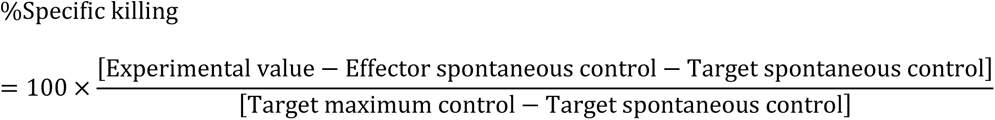

### IL2 secretion assay

I4Z anti-MOG/DR2 BW 5143.4 CAR-T cells were incubated in the presence of DR2+ MGAR cells loaded with different concentrations of MOG or MBP peptide, at different E:T ratios, for 24h. Cells supernatant was collected and tested for IL2 presence, using ELISA assay. Brifely, ELISA plates were incubated in the presence of anti-mouse IL2 ab O.N, followed by blocking for 30 min at room temperature with PBS/2% skim milk. Subsequently, plates were incubated for 2h RT In the presence of cell culture supernatant. After washing, plates were added with biotinylated anti-mouse IL2 ab for 1h RT, followed by three washes and incubation with strep conjugated horseradish peroxidase for 30min RT. Detection was performed using TMB reagent (Sigma). Commercial mouse IL2 was used as positive control and calibration curve.

### Microscopy

Cells were treated as described above for flow cytometry analysis. Z sectioning images with Z-slice of 32.5 μm in 0.5 μm intervals were taken with laser scanning confocal microscope (LSM 700, Zeiss, Germany).

### Mice

DR*1501-Tg mice (a gift from Prof. Vandenbark Portland, Oregon) were bred in-house at the Veterinary Unit, Technion and used at 8-12 weeks of age. All procedures were approved and performed according to institutional guidelines.

### Induction of EAE in DR2-Tg mice

HLA-DR2 mice were screened by FACS for the expression of the HLA transgenes. HLA-DR2 positive male mice between 8 and 12 weeks of age were immunized s.c. at two sites on the flanks with 0.2 ml of an emulsion of 100-200 μg immunogenic peptide and complete Freund’s adjuvant containing 400 μg of heat-killed Mycobacterium tuberculosis H37RA (Difco, Detroit, MI). In addition, mice were given pertussis toxin (Ptx) from List Biological Laboratories (Campbell, CA) on days 0 and 2 post-immunization (2* 200 ng per mouse). Immunized mice were assessed daily for clinical signs of EAE on a 5-point scale: 0 - Clinically normal, 1 - Decreased tail tone or weak tail only, 2 - Hind limb weakness (paraparesis), 3 - Hind limb paralysis (paraplegia), 4 - Weakness of front limbs with paraparesis or paraplegia (quadriparesis), 5 - Paralysis of all limbs (quadriplegia). Mean EAE scores and standard deviations for mice grouped according to initiation of TCRL or vehicle treatment were calculated for each day and summed for the entire experiment (Cumulative Disease Index, CDI, represents total disease load). Daily mean scores were analysed by two-way ANOVA test for nonparametric comparisons between controls and treated groups.

### Recall proliferation

Splenocytes from immunized mice were isolated as described in 8. When indicated, CD4 T cells were isolated using Easy-sep mouse CD4 cell isolation kit (STEMCELL). Cells were than cultured in triplicates with or without peptide antigens for 4 days in a stimulation media (DMEM supplemented with 10% FBS, 2 mM sodium pyruvate, 2 mM l-glutamate, 4 mM 2-mercaptoethanol, and 50 μg/ml penicillin/streptomycin), of which the last 16 h were in the presence of 3H-thymidine to assess proliferation responses. For CD4 T cells cultures, irradiated (3000 rad) splenocytes from naïve mice were added in a 1:2 ratio. Cultures were harvested on glass fiber filters and uptake of 3H- thymidine was assessed by liquid scintillation. The stimulation index (SI) was calculated by dividing the cpm of peptide-stimulated cultures by the cpm of control cultures. The statistical significance of the differences between stimulation levels observed in the different treatment groups was determined Data was compared using Kruskal-Wallis followed by a Dunn’s multiple comparison test.

### CFSE

Isolated splenocytes from immunized mice were stained with CellTrace™ CFSE Cell Proliferation Kit (Thermo) and cultured with or without peptide with different concentrations of 2G10 Ab. After 72 hr CD4+ cell proliferation was evaluated by FACS.

### Statistical analysis

We performed nonparametric Mann–Whitney test or Kruskal-Wallis followed by a Dunn’s test for multiple comparison. EAE score, inhibition and cytotoxicity assays were analysed using 2-way ANOVA test. ****P < 0.0001, ***P < 0.001, **P < 0.01, *P < 0.05.

## Acknowledgments

The authors are grateful to Profs. Arthur A Vandenbark (Neuroimmunology Research and Multiple Sclerosis Research Laboratory, Oregon Health & Science University, Portland, USA) for the HLA-DR2 Transgenic mice and scientific advice and discussions and Lior Mayo (Tel Aviv University, Israel) for scientific advice and discussions.

## Funding

This work was supported by a grant from the Israel Science Foundation (ISF-1889/20) to Y. Reiter.

## Notes

### Competing Interest Statement

The authors have declared no competing interest.

